# Compartment-specific core microbiomes in potato tubers are associated with plant health across genotypes, soils, and years

**DOI:** 10.64898/2026.01.16.699836

**Authors:** Yizhu Qiao, Junqing Qiao, Roeland L. Berendsen, Xu Cheng, Corné M.J. Pieterse, Yang Song

## Abstract

Harnessing plant microbiomes for sustainable agriculture requires understanding not only whether they can boost crop performance, but also how they assemble, persist, and support plant growth and health across environments. While we previously showed that seed tuber microbiomes can predict potato vigour using machine learning, it remained unclear how ecological processes shape tuber microbiome stability and functionality across host genotypes, tuber compartments, soil types, and years. Here, we analyzed the national-scale dataset of 240 field-collected potato seedlots, spanning six genotypes, two soil types, and two growing years, with a focus on the spatially distinct heel and eye compartments of the potato tuber. By profiling over 1,200 bacterial and fungal communities and linking microbiome composition to plant performance using leaf area as a health proxy, we show that plant genotype and tuber compartment are the strongest determinants of microbial diversity and composition. Compartment-specific enrichment of functional traits was observed, with organic compound conversion and nitrogen cycling dominant in the heel, and energy metabolism enriched in the eye. Using a macroecological abundance–occupancy framework, we identified a stable core microbiome of bacterial and fungal taxa that persisted across environments and years. Core members were more strongly associated with plant performance than non-core taxa, and pathogen-suppressive functions were spatially structured, with distinct protective taxa dominating in the heel versus the eye. Together, our findings demonstrate that tuber compartments act as selective microbial filters, shaping persistent microbiomes with specialized functions. By providing an ecological and functional framework for compartment-resolved, stable core microbiomes, this study complements our predictive modelling work and highlights persistent microbial partners as promising targets for improving potato resilience and productivity.

## Introduction

To address the increasing challenges posed by rising food demand, soil erosion, and climate change, the development of sustainable agricultural management strategies is critical (Busby et al. 2017; Compant et al. 2019; Trivedi et al. 2020). One promising solution lies in leveraging the plant microbiome, communities of microbes that live in and around plants, to improve crop productivity and resilience (Berendsen et al. 2012; Haney et al. 2015). These microbial communities can suppress pathogens, mobilize nutrients, and support plant growth under stress (Raaijmakers et al. 2009; Teixeira et al. 2019). However, our ability to harness this potential is limited by an incomplete understanding of how plant-associated microbiomes assemble, persist, and function across diverse environments. A key step toward microbiome-informed crop management is identifying the “core microbiome”, microbial taxa that persistently associate with a host plant across diverse environments and over time, playing a critical role in the plant’s health and productivity. While the concept of a core microbiome has been explored in various crops, critical knowledge gaps remain. In particular, we lack an integrated understanding of how multiple interacting factors, such as plant genotype, specific plant tissue niches, and soil environment, jointly influence microbiome composition, stability, and function. These gaps are especially pressing in the case of potato (*Solanum tuberosum*), the world’s third most consumed crop and a vital food and industrial resource (Devaux et al. 2014).

The growth potential of potatoes, a critical determinant of yield and quality, is influenced by both host factors (e.g., genotype) and environmental factors (e.g., soil type), which shape physiological traits and microbial colonization patterns (Zarzyńska et al.2022; Song et al. 2025). Notably, potato tubers harbor microbial communities that are highly compartmentalized. The heel end (hereafter “heel”), the basal end of the tuber, where it was attached to the stolon, the stem-like structure connecting it to the parent plant, and the eye, the dormant bud that gives rise to new shoots, represent distinct microhabitats that may differentially recruit and maintain microbial taxa (Song et al. 2024, 2025). These spatial niches not only shape microbiome assembly but may also influence key agronomic traits such as sprouting vigor and disease susceptibility. Previous studies have demonstrated that the microbiota of seed tubers can affect crop performance (Sessitsch et al. 2024; Song et al. 2025), but a comprehensive, compartment-resolved understanding of microbiome dynamics across environmental gradients and genotypes is still lacking. Moreover, the mechanisms underlying microbiome assembly and function, as well as their role in maintaining plant health, are still poorly understood.

Although plant-associated microbiomes are critical for various ecological functions, not all microbial members within these communities contribute equally. The core microbiome is typically defined as taxa or functional traits that persistently appear across microbial communities in different habitats (Toju et al. 2018). Here, the persistence of the core microbiome refers to the ability of specific populations to maintain their presence across different habitats over time (Shade & Handelsman 2012; Wagner et al. 2014). These core communities are believed to provide essential and stable ecological functions, making them more easily identifiable across datasets spanning different spatial and temporal scales (Astudillo-García et al. 2017). Specifically, by analyzing multiple possible host-related environmental dimensions, ecological traits such as persistence can highlight which microbiome members are capable of long-term interactions with specific plants (Pfeiffer et al. 2017; Kokou et al. 2019). Given the diversity and complexity of plant microbiomes at multiple scales, identifying core taxa is a crucial step toward understanding their roles in promoting, maintaining, or inhibiting plant and soil health, which is particularly important in potato cropping systems.

This study is the third in our systematic investigation of the potato tuber microbiome. In our first study, we revealed that seed tuber microbiota are vertically transmitted to sprouts and have long-lasting effect on daughter tuber microbiome, highlighting the overlooked importance of the initial microbiome in planting material (Song et al. 2024). In our second study, focusing on the tuber eye microbiome - a subset of the dataset presented here - we demonstrated that seed tuber microbiomes can predict potato vigour, identifying variety-specific microbial indicators through integrating microbiome profiling and drone-based phenotyping (Song et al. 2025). Building on these findings, the present study broadens the scope by analyzing the complete national-scale dataset spanning six genotypes, two soil types, and two growing years. Here, we move beyond vigour prediction to compare the heel and eye compartments of potato tubers, assess the temporal stability of microbiome composition, and identify stable core microbiomes associated with plant health. In doing so, we provide an ecological and functional framework that complements our earlier predictive modelling work and highlights persistent microbial partners as targets for improving potato resilience and productivity.

## Material and Methods

### National-scale potato seed tuber microbiome dataset

We followed the experimental design previously described in the study of Song et al. (2025). Specifically, six genetically divergent potato varieties (from the Royal HZPC Group and Averis Seeds B.V.): Challenger (CH), Colomba (CO), Festien (FE), Innovator (IN), Sagitta (SA), and Seresta (SE) were cultivated in clay soil. To specifically assess the impact of soil type, varieties FE and SE were additionally cultivated in sand soil.

Sample collection and processing followed the procedures described by Song et al. (2025). The same set of samples was used in the present study. In brief, seed tubers from 240 distinct seedlots were sampled over two years. In year 1, 60 seedlots were selected for microbiome analysis of the heel and eye compartments, with four replicate samples per seedlot. Based on the observed consistency among replicates, year 2 sampling included all 180 seedlots with two replicates per seedlot. In total, 600 seedlots were collected. Tissue cores of these 600 potato seedlots were taken using a sterile 0.6 cm corer, pooled, snap-frozen in liquid nitrogen, freeze-dried, and stored at −20°C until further analysis. Freeze-dried tissue samples were placed in 50-mL Falcon tubes with four 5-mm sterile metal beads and homogenized using a heavy-duty paint shaker for 9 min.

### DNA extractions, sequencing and microbial community analysis

Amplicon sequencing and data processing were the same as previously described by Song et al. (2025), and this study used the same sequencing dataset. Briefly, DNA was extracted and libraries were prepared following standard protocols. Bacterial and fungal communities were amplified using primer sets 341F/806R and fITS7/ITS4-Rev, respectively, and sequenced on an Illumina MiSeq platform. Sequencing reads were denoised, merged, and clustered into amplicon sequence variants (ASVs) using the DADA2 pipeline in QIIME2 (v.2023.5) (Callahan et al. 2016; Bolyen et al. 2019). ASVs with fewer than 30 reads or present in fewer than 3 samples were removed. Taxonomic classification was performed using pre-trained classifiers based on the SILVA 99% database (Quast et al. 2012) for bacteria and UNITE v8.0 (Abarenkov et al. 2010) for fungi. Non-target sequences (e.g., mitochondria, chloroplasts) were filtered out. Data were rarefied to 8,000 reads for bacteria and 4,000 for fungi. Alpha diversity was calculated in QIIME2, and Bray-Curtis and weighted UniFrac distance matrices were visualized in R (ggplot2). Community dissimilarities were tested using PERMANOVA (vegan).

The Functional Annotation of Prokaryotic Taxa v.1.0 (FAPROTAX) pipeline was used to extrapolate bacterial community functions. FAPROTAX was constructed by integrating multiple culturable prokaryotic bacteria with reported functions and contained more than 7600 functional annotations for more than 4600 species. This makes it a powerful tool to perform functional annotation based on published metabolic and ecological functions such as nutrient (e.g., C, N, P, and S) cycling, plant pathogens, and symbionts. Given that fungi typically include several guilds (e.g., saprotrophs, pathogens, and symbionts), we further parsed fungal ASVs into these trophic modes based on their taxonomic assignments using the FUNGuild tool (Nguyen et al. 2016). Following the recommendation, we only kept those fungal ASVs with mode assignments that are “highly probable” and “probable’, so as not to overinterpret our results ecologically. In this study, we defined pathogens not only based on those identified within FUNGuild, but also by including common pathogenic fungi frequently reported in the literature.

### Statistical analysis

Abundance-occupancy analysis to detect a core microbiome, Sloan neutral model fit, and data visualization were performed in R (Qiao et al. 2024a). Random forest analysis was conducted to identify the most important ASVs (with higher value of percentage increase in MSE) for leaf area using the “randomForest” R package. The linear discriminant analysis (LDA) effect size (LEfSe) was applied (Wilcoxon P-value□<□0.05, logarithmic LDA score□>□2) to identify the biomarker taxonomy for the different compartments. Nonparametric statistical test (Kruskal-Wallis) was performed to evaluate the alpha-diversity difference and the taxonomical difference among different compartments and years.

## Results

### Diversity and composition of bacterial and fungal microbiomes in potato heel and eye compartments

In our previous study, we used amplicon sequencing to characterize the microbiome of potato tuber eyes and demonstrated that seed tuber microbiomes can predict potato vigour, while also showing that potato genotype and soil type significantly influenced community composition. Here, we reanalyzed the same sequencing dataset to broaden the scope beyond vigour prediction, with a specific focus on the role of tuber compartment (heel and eye) in shaping bacterial and fungal community composition, and on how this compartment effect interacts with potato genotype and soil type. In this dataset, samples were rarified to 8,000 reads for 16S rRNA and to 4,000 for ITS. Shannon diversity analysis revealed that both bacterial and fungal diversity was significantly higher in the heel than in the eye compartment of potato tubers (Fig. 1c, h; *P* < 0.001). To identify distinct microbiome signatures between heel and eye compartment influenced by potato genotype and soil type, we calculated Bray-Curtis dissimilarities for all pairwise sample comparisons and conducted principal coordinate analysis (PCoA). PERMANOVA analysis showed that plant compartment, potato genotype, and soil type all significantly affected both bacterial (Fig. 1d-f) and fungal (Fig. 1i-k) community composition (*P* < 0.001), with potato genotype explaining the largest proportion of variance (24.7% for both bacteria and fungi), followed by plant compartment (bacteria 8.7%, fungi 8.6%), and soil type contributing minimally (bacteria 3.7%, fungi 2.9%). Additionally, PCoA indicated that plant compartment explained the largest variation in microbial communities (5.2-28.6%) across most plant genotypes (Fig. S1).

**Fig. 1.**
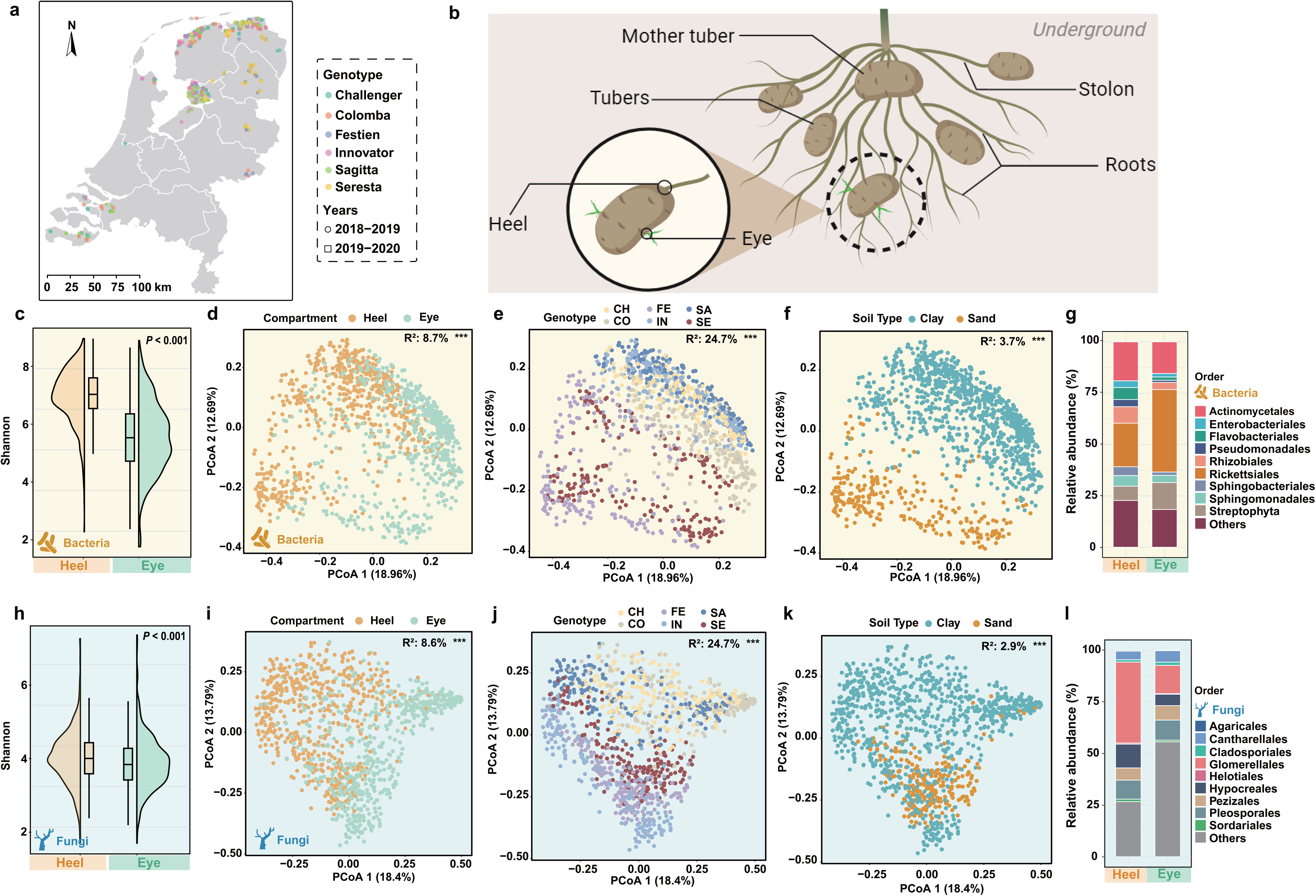
Diversity and composition of bacterial and fungal communities in potato tuber heel and eye compartments. (a) Locations of the 240 fields where the seedlots of six potato varieties were collected in the Netherlands. (b) The different compartments of the potato tuber, focusing on the “eye” and the “heel”. (c) Shannon diversity index of bacterial communities in the eye and heel compartments. (d-f) PCoA of bacterial community composition by compartment (d), genotype (e) and soil type (f). (g) Relative abundance of bacterial orders in the eye and heel compartments. (h) Shannon diversity index of fungal communities in the eye and heel compartments. (i-k) PCoA of fungal community composition by compartment (i), genotype (j) and soil type (k). (l) Relative abundance of fungal orders in the eye and heel compartments.

Microbial communities in potato tuber compartments exhibited distinct taxonomic patterns. Bacterial orders such as Rickettsiales and Actinomycetales were dominant, with Rhizobiales, Flavobacteriales, and Pseudomonadales more enriched in the heel, while Rickettsiales were more abundant in the eye (Fig. 1g). LDA effect size analysis further revealed compartment-specific bacterial genera (Fig. S2a). Similarly, fungal communities showed clear compartmental differences, with Glomerellales (Fig. 1l), *Plectosphaerella*, *Fusarium*, and *Colletotrichum* enriched in the heel, whereas *Cladosporium* and *Stemphylium* were predominant in the eye (Fig. S2b). Overall, while the influence of potato genotype and soil type on tuber microbiomes has been reported previously, our analysis demonstrates that tuber compartment adds a distinct and significant layer of variation. These results highlight clear differences in microbial diversity and community composition between compartments, with compartment and genotype emerging as the primary drivers of microbiome structure.

### Potato genotype-dependent variation in heel and eye microbiomes

Given the significant influence of potato genotype on microbial community composition reported in the previous study, here we specifically investigated how these genotypes effects manifest differently across tuber compartments. Our analysis revealed distinct compartment-specific microbial signatures within each genotype. In the heel, dominant genera included *Flavobacterium*, *Pseudarthrobacter*, and *Streptomyces* (Fig. 2a), while in the eye, *Nocardioides*, *Pseudarthrobacter*, and *Streptomyces* were most abundant (Fig. 2b).

**Fig. 2.**
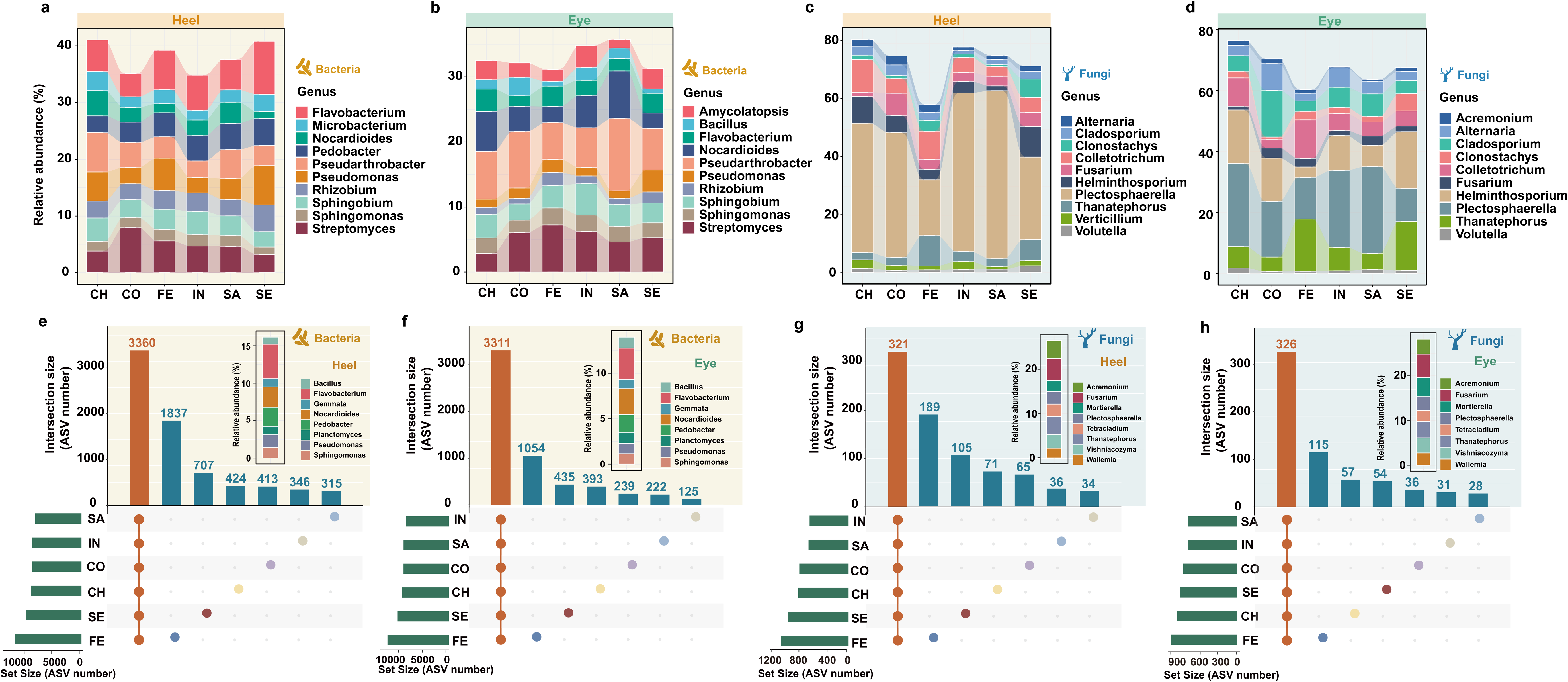
Microbial community composition in heel and eye compartments across potato genotypes. (a-b) Bacterial community composition of tuber heels and eyes of different genotypes. (c-d) Fungal community composition of tuber heels and eyes of different genotypes. (e-f) ASV upset plots of bacterial communities of tuber heels and eyes of different genotypes. (g-h) ASV upset plots of fungal communities of tuber heels and eyes of different genotypes.

We next analyzed the intersection and abundance of ASVs in the heel and eye compartments across genotypes. In the heel, 3,360 ASVs (17% of 20,119 total ASVs) were shared among all genotypes, with *Flavobacterium*, *Nocardioides*, and *Pedobacter* as major contributors (Fig. 2e). Similarly, 3,311 ASVs (19% of 17,874 total ASVs) were shared in the eye, where *Flavobacterium* and *Nocardioides* were dominant (Fig. 2f). The genotype FE exhibited the highest number of unique ASVs in both compartments, 1,837 in the heel and 1,054 in the eye, indicating a strong contribution of overall bacterial diversity, followed by genotype SE.

In fungal communities, dominant genera in the heel included *Plectosphaerella*, *Helminthosporium*, and *Colletotrichum* (Fig. 2c), while the eye was dominated by *Plectosphaerella*, *Helminthosporium*, and *Thanatephorus* (Fig. 2d). Across genotypes, 321 ASVs (17% of 1,917 total ASVs) were shared in the heel, with *Acremonium*, *Fusarium*, and *Thanatephorus* as dominant fungal taxa (Fig. 2g). In the eye, 326 ASVs (19% of 1,755 total ASVs) were shared, primarily represented by *Fusarium* and *Mortierella* (Fig. 2h). Again, genotype FE showed the highest number of unique fungal ASVs in both the heel (189 ASVs) and the eye (115 ASVs). In summary, potato genotypes harbored distinct microbial community structures in both the heel and eye compartments. The FE genotype consistently supported greater microbial diversity, and only a small proportion of ASVs (17∼19%) were shared across all genotypes, indicating a high degree of genotypes specificity in microbiome assembly in the heel and eye compartment.

### Effect of soil type on microbial community composition in heel and eye compartments

To investigate how soil type influences microbial community composition in the heel and eye compartments of potato tubers, we analyzed samples from two contrasting soil types: clay and sand. The analysis focused on two potato genotypes, FE and SE, which were grown in both soil types. We assessed differences in community composition using PCoA based on Bray-Curtis and weighted UniFrac distances, which reflect compositional and phylogenetic variation, respectively. PCoA based on Bray-Curtis distances revealed clear separation between the microbial communities from clay and sandy soils in both the heel and eye compartments for both genotypes (Fig. 3a, b). These high R² values indicate that soil type is a major factor shaping microbial community composition in these potato tuber compartments. In contrast, PCoA based on weighted UniFrac distances, accounting for both taxonomic composition and phylogenetic relatedness, showed lower R^2^ values. Specifically, the R² values for the heel microbiomes were 0.14 for both the FE and SE genotypes, while for the tuber eye microbiomes, the R² values were 0.38 (FE) and 0.25 (SE) (Fig. 3a, b). These results suggests that while soil type strongly influences microbial community composition, it has a more limited effect on the overall phylogenetic structure of these communities. In both the heel and eye compartments, different ASVs were selected in sandy versus clay soils. However, these ASVs were closely related phylogenetically, indicating that the observed shifts occurred primarily at the taxonomic level rather than reflecting deep evolutionary divergence.

**Fig. 3.**
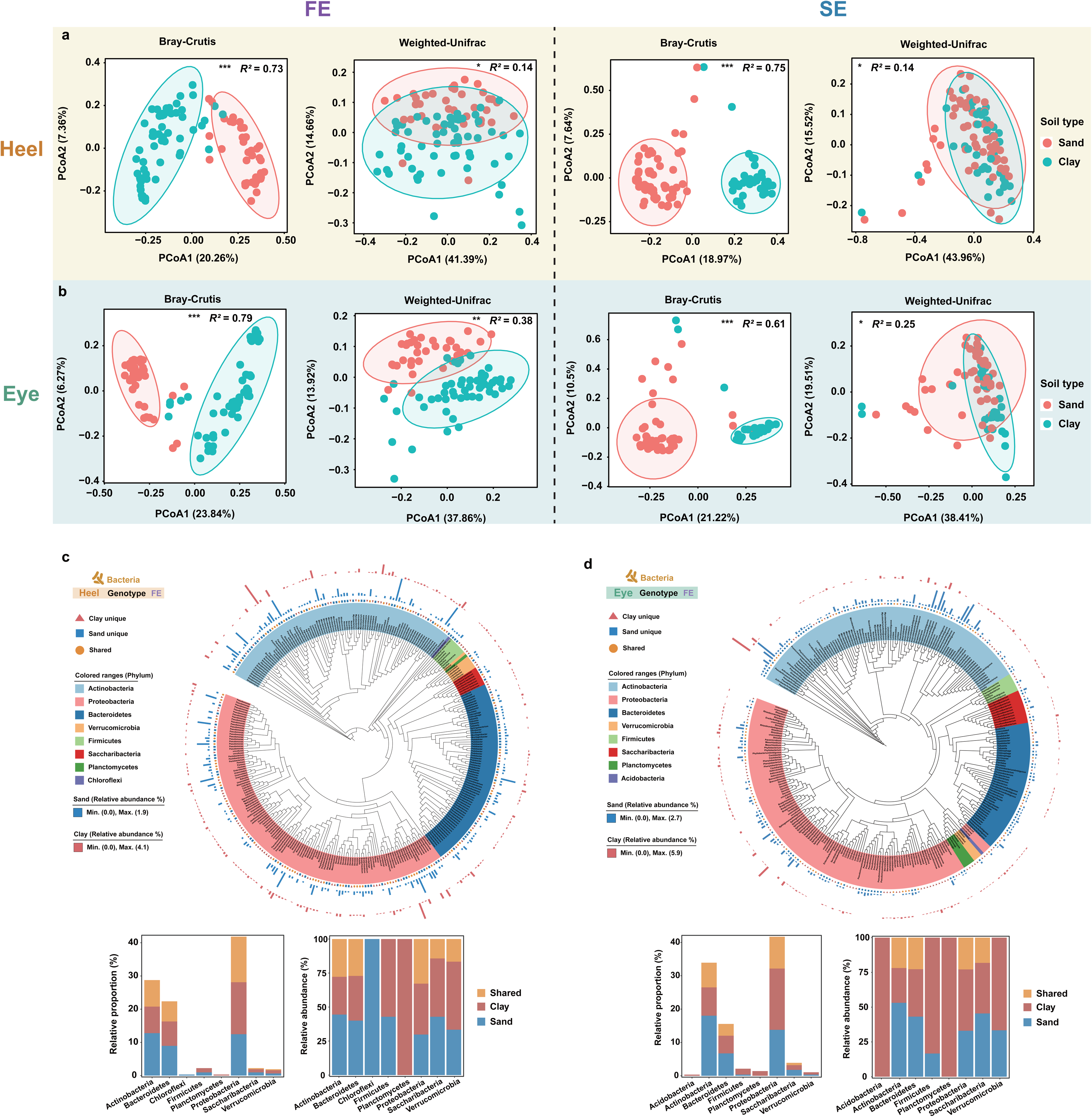
Effect of soil type on microbial community composition in potato heel and eye compartments. (a-b) PCoA of microbial community composition in the heel (a) and eye (b) compartments across different soil types. (c-d) Bacterial phylogenetic tree of tuber heels (c) and eyes (d) in clay and sand soils in FE genotypes. The bar plot quantifies the relative proportion (left) and relative abundances (right) of these specific phylum-level taxa in different soil types.

Interestingly, for other factors such as plant compartment, the R² value derived from UniFrac distances were higher than those from Bray-Curtis (Fig. 1d; Fig. S3a), suggesting that compartment-specific microbial differences may be associated with evolutionary selection pressures. To further explore this, we constructed phylogenetic trees of the high-abundance microbial communities (relative abundance >0.1%) for both compartments and annotated them by source and taxon abundance (Fig. 3c, d; S4; S5). These phylogenetic trees and corresponding bar plots showed that high-abundance genera were evenly distributed across soil types, especially in both heel and eye compartments (Fig. 3c, d; Fig. S5). However, certain genera exhibited notable differences in abundance between soil types, for example, Chloroflexi was more abundant in sandy soils, while Planctomycetes was more enriched in clay soil.

Overall, our results indicate that although soil type is an important environmental factor, microbial responses to soil conditions are primarily realized through the selection of closely related taxa with differential abundances across compartments. This highlights the presence of compartment-specific ecological filtering within the same host.

### Compartment-specific functional profiles and pathogen distributions in the potato tuber microbiome

To better understand the ecological roles and potential risks associated with tuber-associated microbiomes across different compartments, we next examined both the predicted metabolic functions of bacterial communities and the distribution of potential pathogens. In our analysis of predicted bacterial metabolic and ecologically relevant functions, we found that the heel-associated bacterial community showed increased functional potential for xylanolysis (+700%), nitrite ammonification (+137%), ureolysis (+68%), and cellulolysis (+63%) (Fig. 4a). In contrast, the eye-associated bacterial community showed increased functional potential for aerobic ammonia oxidation (+55%), sulfur respiration (+58%), iron respiration (+110%), and respiration of sulfur compounds (+61%) (Fig. 4a).

**Fig. 4.**
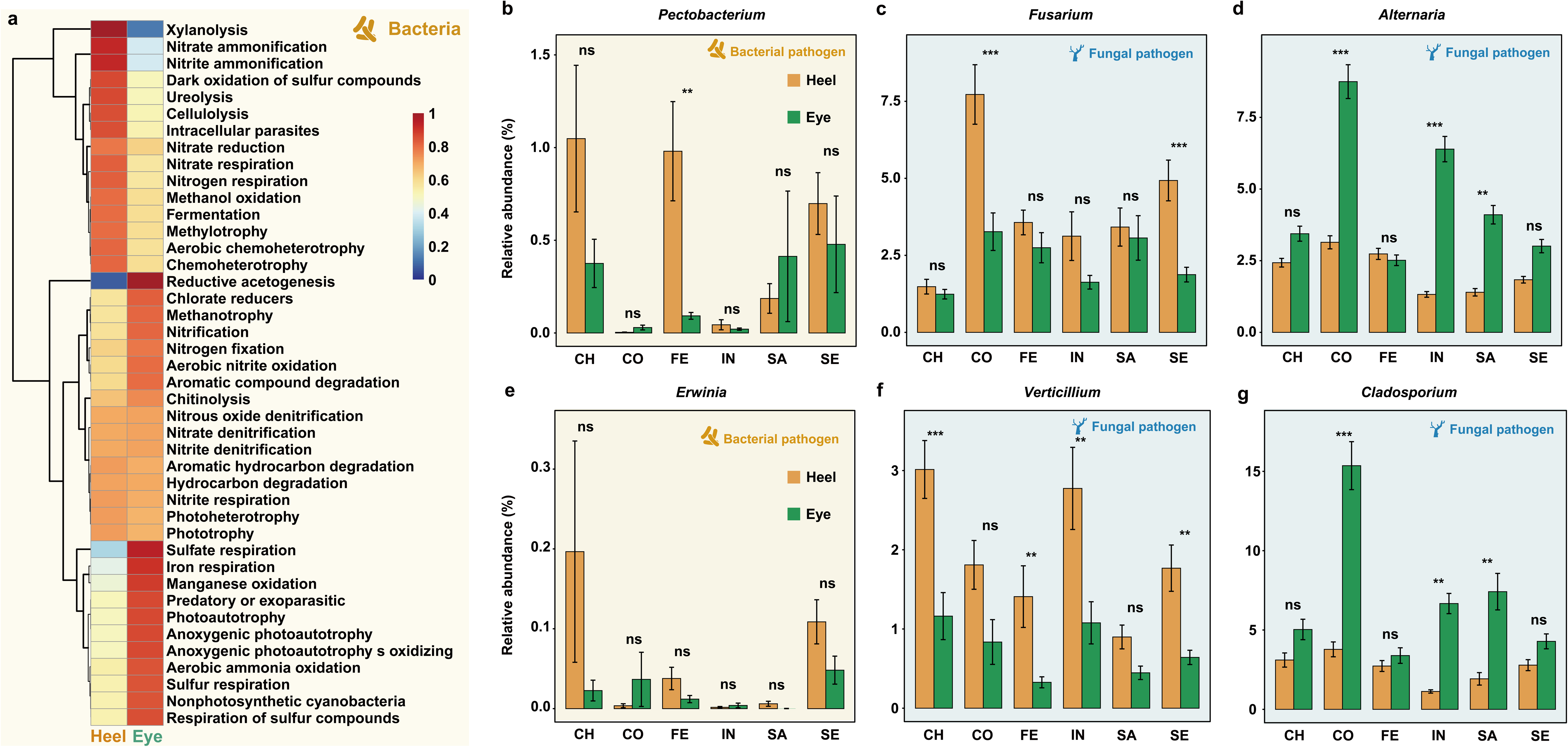
Differential functional traits of bacteria and fungi in tuber heel and eye compartments. (a) FAPROTAX annotated differences in functional profile in different compartments (heel and eye) of potato. (b-g) Relative abundance of different pathogenic bacteria (both bacteria and fungi) on different compartments (heel and eye) of potato. Pathogen included *Pectobacterium*, *Erwinia* (bacterial pathogens) and *Fusarium*, *Alternaria*, *Verticillium*, *Cladosporium* (fungal pathogens).

To assess pathogen distribution, we defined pathogens by integrating assignments from FUNGuild, a database-based tool that annotates fungal taxa according to their ecological guilds, together with bacterial and fungal taxa commonly reported in the literature as pathogenic. We then analyzed the relative abundance of bacterial and fungal pathogens in the heel and eye compartments across potato genotypes. In most genotypes (except FE), the abundance of bacterial pathogens (*Pectobacterium* and *Erwinia*) was slightly higher in the heel compared to the eye, although the differences were not statistically significant in each genotype (Fig. 4b, e). In contrast, fungal pathogens such as *Fusarium* and *Verticillium* were significantly more abundant in the heel than in the eye in most genotypes (except SA), with particularly strong differences observed in the SE genotype (*P* < 0.01) (Fig. 4c, f). Conversely, *Alternaria* and *Cladosporium* were significantly more abundant in the eye compartment compared to the heel in specific genotypes, especially CO, SA and IN (*P* <0.01) (Fig. 4d, g). These findings highlight distinct compartment-specific patterns in both functional profiles of bacterial communities and the abundance of potential pathogens, suggesting spatial differentiation in microbial roles and risks within the potato tuber.

### Stable core microbiomes in potato tuber compartments across genotypes, soils, and years

To determine whether different compartments of potato tubers harbor stable microbial assemblages across genotypes, soil types, and years, we focused on microbial taxa that maintained the most stable and persistent associations with the host. We observed numerous bacterial ASVs shared across all genotypes, soil types, and years in both the heel and eye compartments (Fig. 2e-h; Fig. S3; Fig. S6), with a smaller but still notable overlap in the fungal communities. These findings suggested that, despite pronounced differences among genotypes and strong biogeographic signals, both heel and eye compartments consistently recruits many of the same microbial taxa, which may be functionally important for the host.

To specifically assess the stability of microbial associations at the compartment level, we explored abundance-occupancy distributions of taxa to infer the bacterial and fungal core microbiomes in the heel and eye compartments. Core taxa were defined as those with an occupancy greater than 0.9 (i.e., taxa present in at least 90% of samples across genotypes, soil types, and years; Fig. 5a-d). Among bacteria, 18 taxa (22 ASVs) were cosmopolitan in the heel compartment (Fig. 5e), including several abundant Proteobacteria, with *Sphingobium* as the dominant genus (mean relative abundance: 2.49%), and Actinobacteria, with *Pseudarthrobacter* as the most abundant genus (mean relative abundance: 4.28%). The heel core bacterial microbiome also included genera with potential plant beneficial functions (e.g., *Sphingomonas*, *Rhizobium*, *Streptomyces*) and several associated with disease suppression (e.g., *Sphingopyxis*). Notably, genera such as *Rhizobium*, *Rhizobiaceae*, and *Phyllobacterium* were core members exclusive to the heel compartment.

**Fig. 5.**
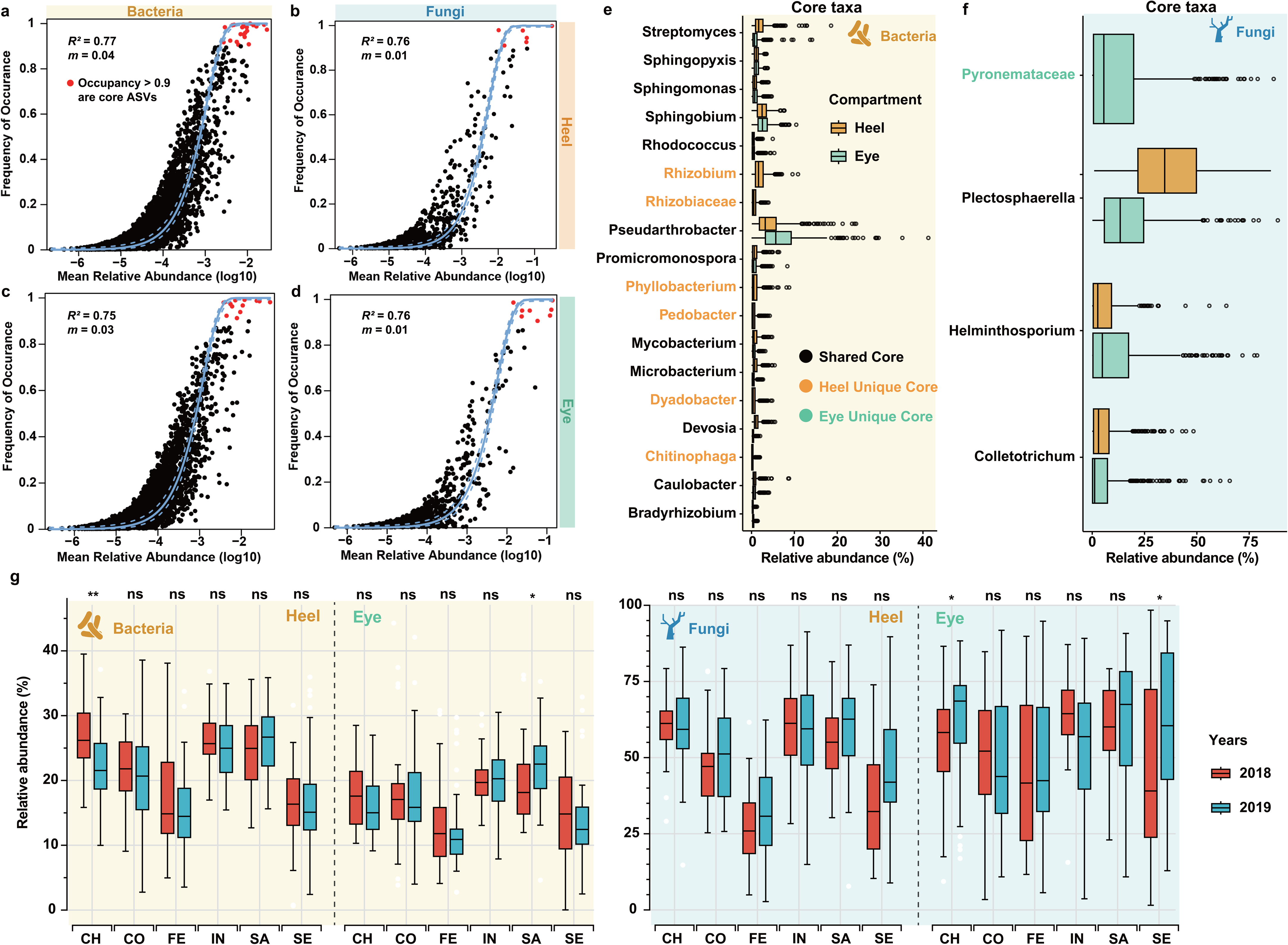
Microbial prevalence in heel and eye compartments of potato tubers. **(a-d)** Abundance-occupancy distributions were used to identify core members of the heel (a-b) and eye (c-d) microbiome for bacteria and fungi. The solid line represents the fit of the neutral model, and the dashed line is 95% confidence around the model prediction. Taxa with an occupancy > 0.9 (i.e., detected in all samples) were considered members of the core. (e-f) Relative abundance of these bacterial (e) and fungal (f) core taxa is represented as boxplots. Taxa exclusive to a genotype are indicated in orange (heel) or green (eye), and taxa shared across both genotypes are black. (g) The combined relative abundance of all core taxes in 2018 year (red) and 2018 year (blue). Stars above box plots represent statistical significances as determined by Wilcoxon test (***: *P* ≤ 0.001, **: *P* ≤ 0.01, *: *P* ≤ 0.05).

In the eye compartment, 12 bacterial taxa (13 ASVs) were identified as core members (Fig. 5d), again dominated by Proteobacteria and Actinobacteria, with *Pseudarthrobacter* (mean relative abundance: 7.21%) and *Sphingobium* (mean relative abundance: 2.71%) as the most abundant genera.

For fungi, 3 core taxa (5 ASVs) were identified in the heel, all belonging to the phylum Ascomycota, with *Plectosphaerella* as the dominant genus (mean relative abundance: 35.8%) (Fig. 5f). In the eye compartment, 4 core fungal taxa (8 ASVs) were identified, also from Ascomycota, again dominated by *Plectosphaerella* (mean relative abundance: 17.6%) (Fig. 5f), with *Pyronemataceae* emerging as a core fungal genus unique to this compartment.

To assess the temporal stability of core taxa, we compared their presence across the two sampling years. The core taxa were consistently detected and showed no significant differences between years (Fig. 5g), indicating their persistence over time. Together, these findings suggest that the heel and eye compartments of potato consistently harbor distinct, yet stable, core microbiomes that persist across diverse host genotypes, environmental conditions, and temporal scales.

### Impact of core microbiomes on plant health

To explore the potential impact of core microbiomes on plant health, we reanalyzed the potato leaf area dataset from our previous study in combination with the core microbiome profiles identified here. Because leaf area can vary across years and genotypes, we standardized it within each year and genotype by z-score transformation to enable cross-sample comparisons (Fig. 6a). The results showed that, in both bacterial and fungal communities, the importance of individual ASVs in predicting leaf area increased with their prevalence (Fig. 6b, c). This suggests that more prevalent ASVs are more strongly associated with leaf area, implying a potential role in promoting plant growth. The consistent trend across both microbial domains highlights microbial prevalence as a key factor in predicting plant functional traits such biomass and vigor. Notably, in both heel and eye compartments, core microbial groups contributed more to leaf area prediction than non-core groups (Fig. 6d, e), suggesting that core taxa have a stronger influence on plant performance than transient or rare community members.

**Fig. 6.**
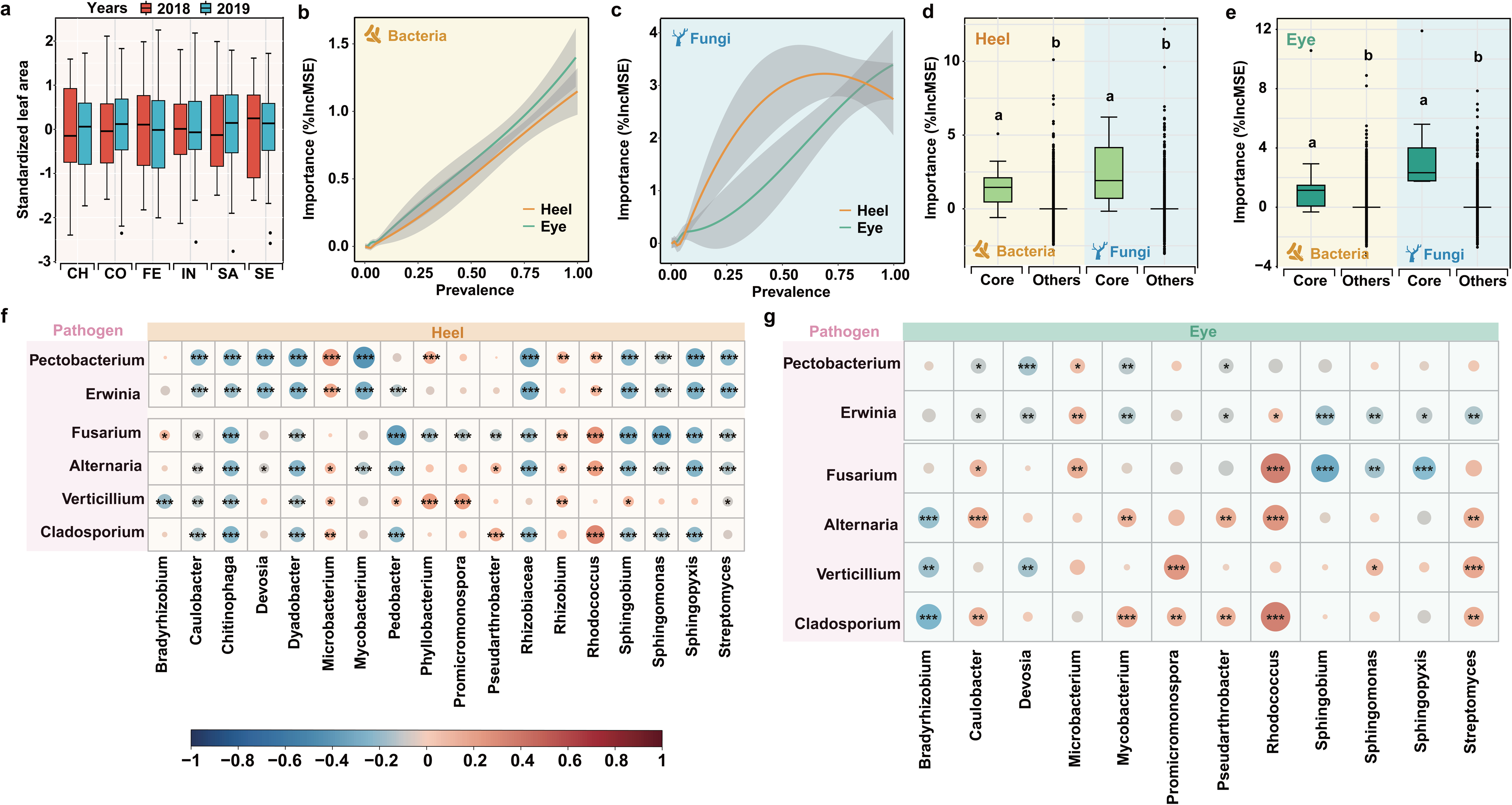
Core microbial communities associated with plant health. (a) The variation in leaf area across different potato genotypes and years. Considering that the leaf area varies across years and genotypes, we standardized leaf area according to each year and genotype. (b-c) The relationship between prevalence and the importance of bacterial (b) and fungal (c) ASVs in predicting leaf area. The shaded regions represent standard errors. (d-e) Boxplots comparing the importance of core (green) and other (purple) bacterial (orange) and fungal (blue) group in predicting leaf area for the heel (d) and eye (e) compartments. Different letters indicate statistically significant differences between groups (p < 0.05). (f-g) The relationships (Spearman correlation) between core taxa and pathogen abundance in the heel (f) and eye (g) compartment, with positive (red) and negative (blue) correlations denoted by color intensity. Significant correlations are marked with asterisks.

To further investigate how core microbiomes influence plant health, we conducted correlation analyses between core microbial taxa and pathogen abundance. Several core bacterial genera, including *Chitinophaga*, *Dyadobacter*, and *Streptomyces*, exhibited significant negative correlations with known plant pathogens, including *Fusarium*, *Alternaria*, and *Verticillium* (Fig. 6f). Interestingly, the nature of these interactions varied between compartments: in the heel, bacterial taxa displayed stronger overall negative correlations with pathogen abundance, whereas in the eye, specific taxa such as *Bradyrhizobium*, *Sphingobium* and *Sphingomonas* played more dominant roles (Fig. 6g). This compartment-specific pattern highlights the niche-dependent functionality of microbiome members and suggests spatial variation in microbial contributions to pathogen suppression. Collectively, these findings indicate that core microbiome members in different compartments are not only stable across genotypes and environments, but also play an active role in supporting plant health by mitigating pathogen pressure.

## Discussion

Uncovering the ecological principles that shape plant microbiome assembly is essential for understanding host-microbe interactions and applying this knowledge to improve crop productivity (Busby et al. 2017; Berg et al. 2017). In this study, we focused on the differences in microbial communities between the heel and eye compartments of potato tubers. Building on our previous work on genotype, soil type, and temporal variation, we analyzed these factors from a compartmental perspective to uncover compartment-specific ecological filtering and to identify core microbial members consistently linked to potato health.

### Compartmental differentiation of microbial diversity and composition

In our study, bacterial and fungal diversity was consistently higher in the heel than in the eye compartment of potato tubers, consistent with previous findings (Song et al. 2024). This likely reflects heel’s connection to the mother plant and nutrient inputs, making it a reservoir for beneficial microbes (Hu et al. 2016; Berg et al. 2017). In contrast, the eye represents a selective, nutrient reserved niche with reduced diversity, possibly to support dormancy and sprouting (Wei et al. 2021; Song et al. 2024). Our results reveal that the composition of microbial communities is largely determined by plant genotype and plant compartment, echoing findings in maize (Walters et al. 2018) and rice (Zhang et al. 2019). For example, the enrichment of *Flavobacterium* and *Pseudomonas* in the heel aligns with their known roles as decomposers of complex organic material in rhizosphere environments (Mendes et al. 2013). In the eye, the predominance of *Rahnella* and *Gluconacetobacter* suggests metabolic specialization for exploiting its unique nutritional context, similar to microbial strategies observed in germinating seeds (Truyens et al. 2015).

### Heel and eye microbiome variation across potato genotypes

Potato genotypes differentially shaped microbial communities across heel and eye compartments, with both genotype-specific and conserved patterns. This aligns with broader evidence that host traits, such as exudate composition or tissue-specific traits, influence microbiome recruitment (Bulgarelli et al. 2012; Edwards et al. 2015). Despite this variability, a subset of bacterial genera, including *Flavobacterium* and *Nocardioides*, was consistently found across genotypes and compartments, suggesting the presence of a compartment-specific core microbiome linked to essential functions such as phosphorus solubilization, phytohormone production, and pathogen suppression (Kolton et al. 2013; Zhang et al. 2019). Interestingly, the FE genotype exhibited the highest number of unique ASVs, pointing to an elevated capacity for microbial recruitment and community diversification. This supports ecological theory that links microbial diversity to greater functional redundancy and ecosystem resilience (Heijden et al. 2007).

### Soil type drives compartment-specific potato microbiome composition but not phylogeny

Our study demonstrates that soil type exerts a compartment-specific influence on the assembly of potato tuber-associated microbial communities. This aligns with prior findings that soil physicochemical properties, such as texture, nutrient content, pH, and moisture retention, serve as environmental filters that shape microbial colonization (Lauber et al. 2009). Unifrac distances and phylogenetic analyses revealed that dominant microbial lineages were conserved across soil types, indicating that soil-driven selection acts primarily at the abundance level rather than through broad phylogenetic restructuring. Importantly, these effects were not uniform across compartments. In the heel, soil type drove strong compositional shifts at the abundance level (high Bray-Curtis R²) but had little impact on phylogenetic structure (low UniFrac R²), suggesting a buffered and stable ecological niche. By contrast, in the eye, soil type explained higher variation in both taxonomic composition and phylogenetic structure (Bray-Curtis and UniFrac R² values), indicating that the eye microbiome is more dynamic and responsive to soil conditions (Petrushin et al. 2024). This compartment-specific soil type effect highlights that different microhabitats within the same host impose distinct ecological filters: the heel appears to support a conserved and resilient community, whereas the eye acts as a more open and selective interface between host and environment. These findings extend principles established for rhizosphere microbiomes to belowground storage organs, emphasizing that local microhabitat context fundamentally modulates how soil properties shape plant-associated microbial communities.

### Compartment-specific metabolic potentials and pathogen niches

Functional profiling revealed distinct metabolic potentials between compartments. The heel was enriched in carbon and nitrogen cycling functions, suggesting a role in nutrient turnover at the stolon-root interface (Berendsen et al. 2012; Zhalnina et al. 2018). In contrast, the eye compartment showed higher potential for sulfur and iron respiration, suggesting a functional specialization in these redox processes (Shu et al. 2016). These findings highlight a spatial partitioning of microbial functional roles within the tuber, complementing observed taxonomic differences. Pathogen profiling also revealed distinct compartment-specific distribution patterns. Fungal pathogens such as *Fusarium* and *Verticillium* were more abundant in the heel, while airborne fungi like *Cladosporium* were enriched in the eye. These differences likely reflect distinct exposure routes and ecological niches (Zhalnina et al. 2018). Such spatial structuring of pathogen communities underscores the need for compartment-targeted disease management strategies.

### Ecological roles and temporal stability of core microbes

Core microbiomes, defined as taxa consistently associated with hosts across environments, were dominated by Proteobacteria and Actinobacteria in both compartments (Shade & Handelsman 2012; Hernandez-Agreda et al. 2017; Kokou et al. 2019). Core genera such as *Sphingobacterium*, *Sphingomonas*, and *Pseudoarthrobacter* are known for roles in hormone regulation, nutrient uptake and stress resistance (Innerebner et al. 2011; Qiao et al. 2024b), suggesting potential functional contributions to plant health. In fungi, Ascomycota dominated, with core genera such as *Plectosphaerella* potentially functioning either as latent pathogens or as endophytes with immunomodulatory or antimicrobial properties (Zhou et al. 2022). Pyronemataceae, a saprophytic family unique to the eye, may contribute to mineral nutrient uptake (Heijden et al. 2007). The consistent detection of core taxa across two years highlights their ecological persistence and potential functional importance. This temporal stability supports the view of core microbiota as evolutionarily selected partners that help buffer environmental variation (Kokou et al. 2019; Vandenkoornhuyse et al. 2015). By integrating community composition with functional potential and temporal consistency, our study reinforces the value of targeting core microbiomes in sustainable crop management and provides a foundational reference for future microbiome-informed breeding or inoculation strategies.

### Potential functional links between core taxa and plant health

Consistent with previous studies highlighting the link between microbial occupancy and functional impact (Shade & Handelsman 2012; Agler et al. 2016), we found that taxa with higher prevalence contributed more to leaf area prediction, indicating a stronger link to plant performance. Notably, *Sphingomonas* emerged as a key taxon showing strong negative correlations with pathogen abundance across both heel and eye compartments, implying a broad-spectrum protective effect (Innerebner et al. 2011). Similarly, *Chitinophaga*, enriched in the heel and negatively associated with pathogenic taxa, likely exerts antifungal effects through the secretion of chitinases and secondary metabolites such as phenazines and polyketides (Carrión et al. 2019). Moreover, core bacterial taxa in the heel exhibited stronger negative associations with pathogens than those in the eye, indicating compartment-specific ecological roles shaped by local microhabitat conditions. Altogether, our findings reinforce the ecological and functional significance of core microbiomes as not merely stable components of the plant microbiota but as active agents in enhancing plant health through pathogen suppression, nutrient mobilization, and potential immune modulation. These insights underscore the value of targeting core microbiota in future microbiome-based agricultural applications aimed at improving crop resilience and productivity.

## Data availability

The raw sequence data generated from this study are available at https://www.ncbi.nlm.nih.gov/bioproject/PRJNA1091851/.

## Acknowledgements

We acknowledge the funding support received from Europees Landbouwfonds voor Plattelandsontwikkeling (ELFPO) on the “Flight-to-vitality” project. This work was also partly supported by the Horizon Europe research and innovation programme, project BOLERO (HORIZON-CL6-2021-BIODIV-01-13, grant agreement no. 101060393). We gratefully acknowledge financial support from the China Scholarship Council.

## Supplementary information

Additional file 1: Supplementary Figures S1-S6.

