## supplemental file for "Compartment-specific core microbiomes in potato tubers are associated with plant health across genotypes, soils, and years"

*b, Key lab of organic-based fertilizers of China and Jiangsu provincial key lab for solid organic waste utilization, Nanjing Agricultural University, Nanjing 210095, China*

Corresponding author:

* Yang Song,

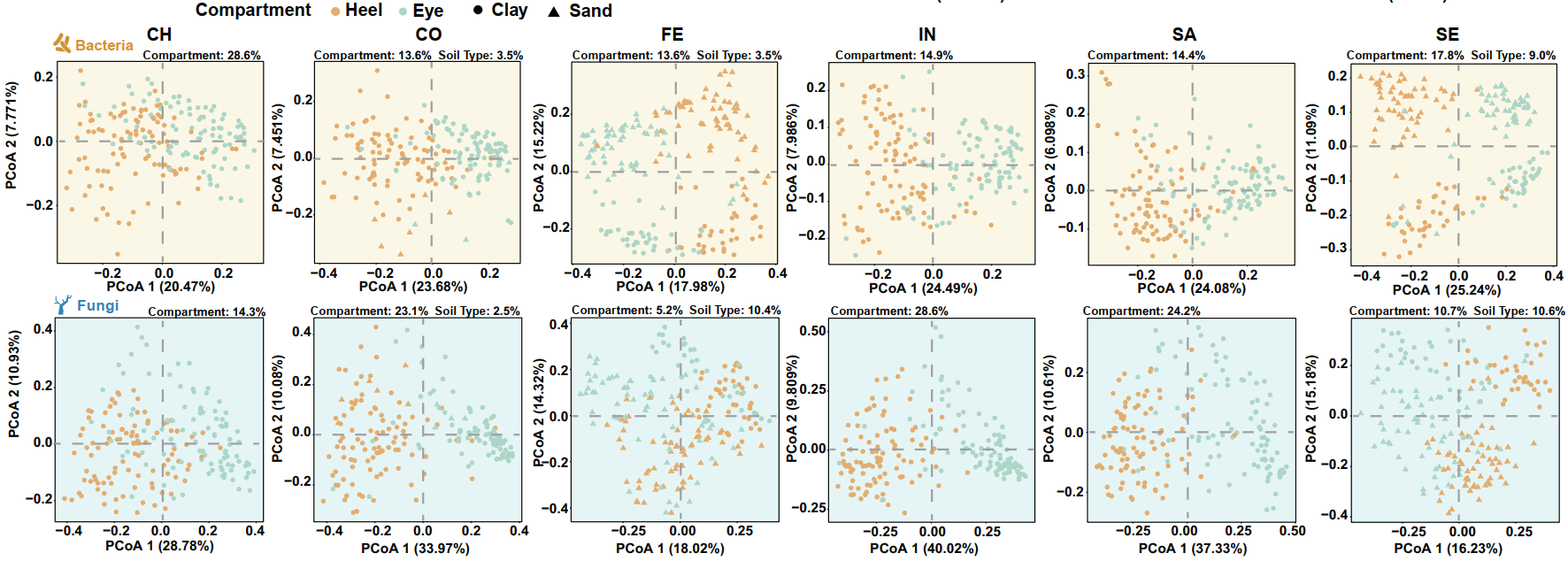

**FIG S1 Principal Coordinate Analysis (PCoA) based on Bray-Curtis distances for microbial communities in different genotypes.**

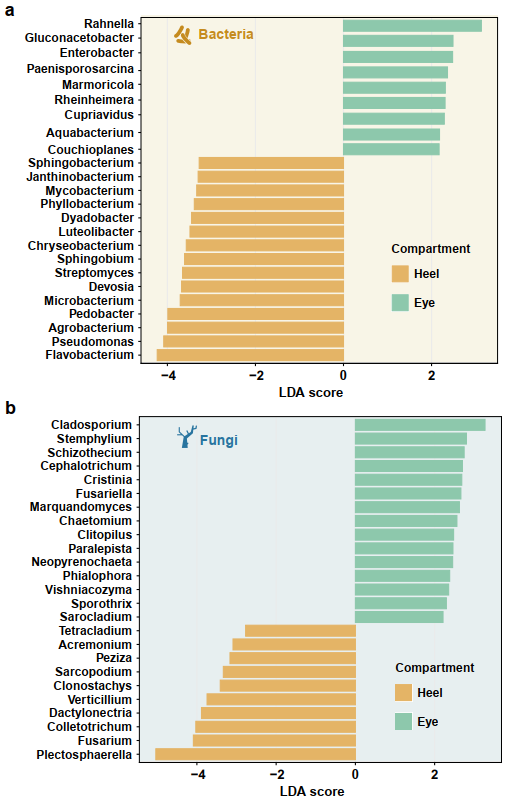

**FIG S2 The linear discriminant analysis effect size (LEfSe) analysis of microbial abundance among different compartments.** (a) Taxa until bacterial genus level with significant differences heel and eye were detected by LEfSe analysis with a LDA threshold score of 2 and a p-value of 0.05. (b) Taxa until fungal genus level with significant differences heel and eye were detected by LEfSe analysis with a LDA threshold score of 2 and a p-value of 0.05.

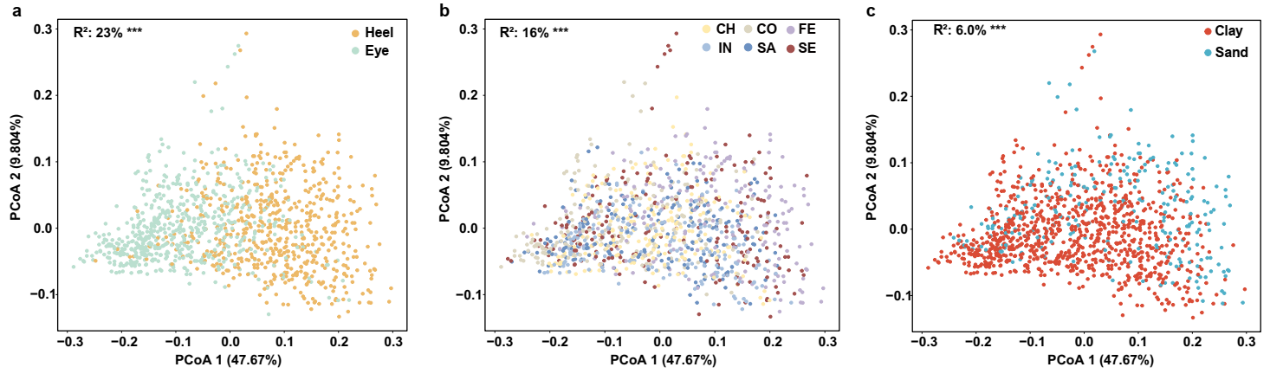

**FIG S3** Based on PCoA analysis (UniFrac distance), the differences in bacterial community structures between the tuber eye and heel (a), genotypes (b) and soil types (c).

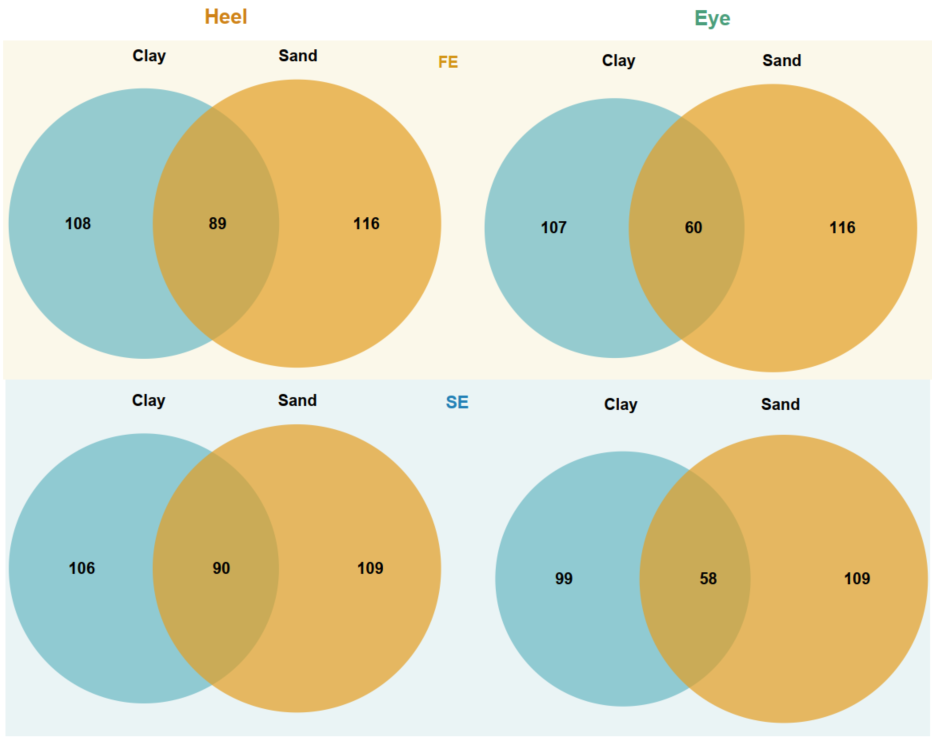

**FIG S4** We constructed the phylogenetic trees of high-abundance (relative abundance > 0.1%) ASVs and plotted the corresponding Venn diagrams of these high-abundance ASVs. The unique and shared ASVs in the Venn diagrams corresponded to the sources of the labelled ASVs on the phylogenetic tree (sand, clay and shared).

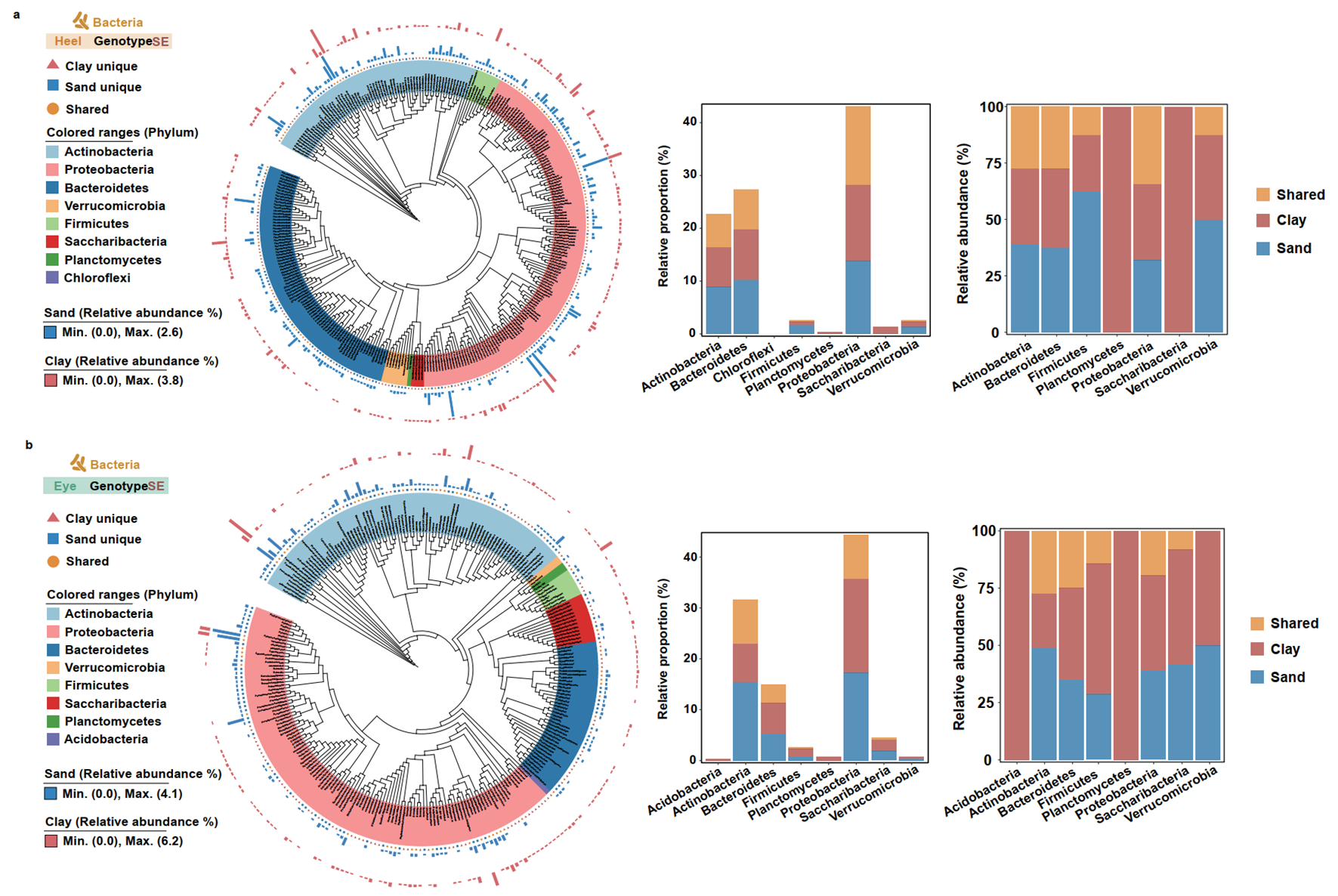

**FIG S5** Bacterial phylogenetic tree of tuber heels (a) and eyes (b) in clay and sand soils in SE genotypes. The bar plot quantifies the relative proportion (left) and relative abundances (right) of these specific phylum-level taxa in different soil types.

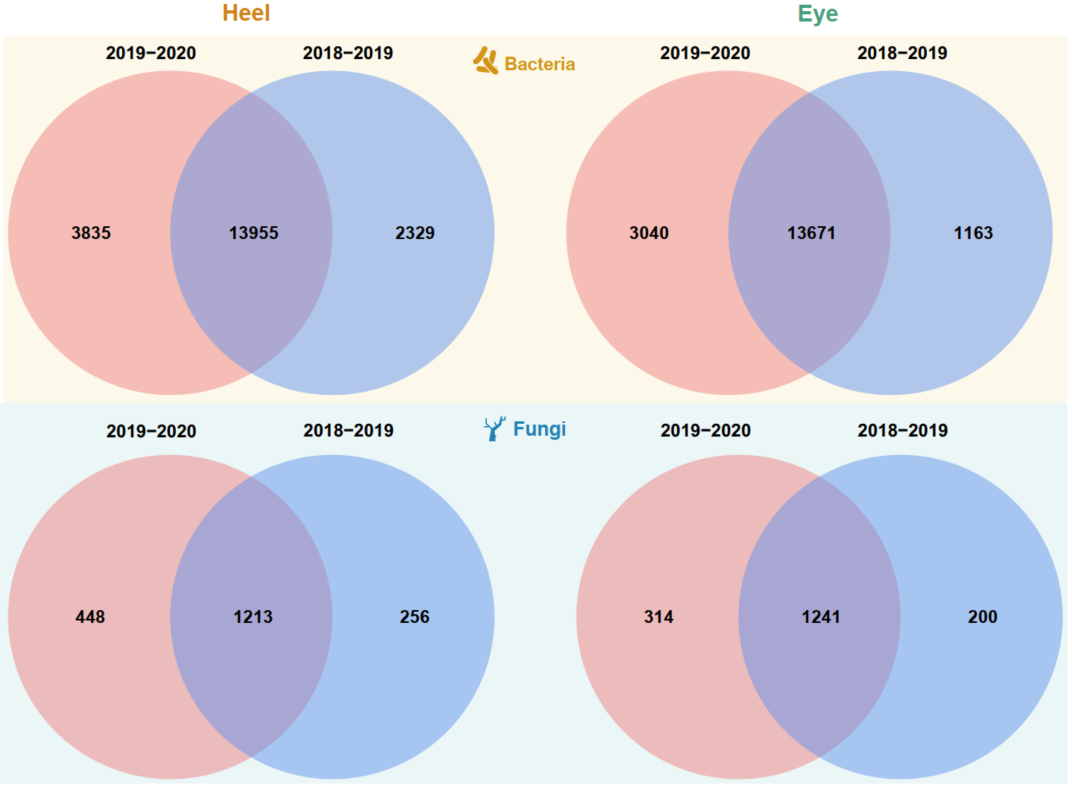

**FIG S6** Venn diagrams illustrating the distribution of bacterial and fungal ASVs between year 1 and 2 in the heel and eye.
